# Prevalence, mechanisms and comparison of detection methods of fosfomycin resistance in *E. coli* from urinary tract infections

**DOI:** 10.1101/234435

**Authors:** Jennifer L. Cottell, Mark A. Webber

## Abstract

As numbers of bacterial isolates resistant to first line antibiotics rise there has been a revival in the use of older drugs such as fosfomycin. Fosfomycin is a cell wall inhibitor with a unique mode of action, increasingly used in the treatment of urinary tract infections. In this study, the prevalence of fosfomycin resistant *E. coli* in a panel of 1000 urine isolates was investigated. Three different clinically used fosfomycin susceptibility testing methods were assessed and genome sequencing used to characterise resistant isolates.

Of the 1000 isolates, 676 were *E. coli* of which initial susceptibility testing with the MAST Uri®system suggested 81 (12%) were fosfomycin resistant. Of these, 62 were subsequently confirmed as being *E. coli.* However, using micro-broth dilution, agar dilution and E-test strips, a lower rate of 1.3% (8/62) of *E. coli* isolates were robustly identified as being truly fosfomycin resistant; a prevalence comparable with other similar studies. The use of E-test and 96-well breakpoint plates gave results that were inconsistent and hard to interpret. Resistant isolates of *E. coli* belonged to diverse MLST types and each had a unique set of chromosomal alterations in genes associated with fosfomycin resistance. Changes in GlpT and UhpT/UhpA transport systems were commonly identified, with 6/8 of the resistant isolates possessing amino-acid changes or deletions absent in susceptible strains. Fosfomycin resistant isolates were not multiply drug resistance and did not carry plasmidic fosfomycin resistance genes. Therefore, the use of fosfomycin may be unlikely to drive selection of a particular clone or movement of transferrable resistance genes.

Fosfomycin remains a viable option for the treatment of *E. coli* in uncomplicated UTIs, different susceptibility testing platforms can give very different results regarding the prevalence of fosfomycin resistance with false positives a potential problem that may unnecessarily limit use of this agent.

## Introduction

Globally, increasing numbers of infections are caused by bacteria resistant to current antibiotics.[1] As there is a lack of new antibiotics in development, the revival of older drugs with distinct methods of action has been proposed as a short-term solution.[2] One such drug is fosfomycin, a cell wall inhibitor discovered in 1969, with a novel mode of action and broad spectrum activity.[3] Fosfomycin is a phosphonic-acid derivative which enters bacterial cells by active transport. In *Enterobacteriaceae*, fosfomycin is taken up by mimicking the natural substrates of two nutrient transport uptake systems, the constitutively functional l-α-glycerol-3-phosphate transporter, GlpT, and the hexose-uptake system, UhpT, which is inducible in the presence of glucose-6-phosphate.[4] Expression and regulation of genes encoding these systems requires the presence of cyclic AMP (cAMP), cAMP-receptor protein complexes and regulatory genes including *uhpA*.[5-7] Once in the bacterial cytosol, fosfomycin acts as a phosphoenolypyruvate analogue preventing the initial step of cell wall synthesis, via inhibition of MurA (enzyme UDP-N-acetylglucosamine enolpyruvyl-transferase).[4] Binding of fosfomycin to a key residue (Cys115 in *E. coli*) in the active site of MurA prevents the formation of UDP-GluNAc-enolpyruvate, which impedes successful N-acetylmuramic acid and peptidoglycan biosynthesis leading to cell death.[8] This unique mechanism of action allows a broad-spectrum of activity against both Gram-positive and Gram-negative bacteria. As fosfomycin acts prior in the biosynthesis pathway to other cell wall inhibitors β-lactams and glycopeptides it is not inhibited by resistance determinants which act against these drugs. Therefore, fosfomycin retains activity against methicillin resistant *Staphylococcus aureus* (MRSA) and *Enterobacteriaceae* producing extended spectrum beta-lactamases (ESBLs).[9]

Several mechanisms have been reported to confer fosfomycin resistance, these include decreased drug uptake, upregulation of MurA, and enzymatic drug inactivation. Historically the most commonly documented mechanism of resistance has been impaired transport of fosfomycin into the cytoplasm, due to mutations in structural or regulatory genes of the nutrient transport systems.[10] In *E. coli*, insertions, deletions or mutations leading to amino-acid changes in *glpT*, *uhpT* or *uhpA* have been associated with impaired uptake of fosfomycin and with reduced susceptibility both *in-vitro* and *in-vivo*. Alternatively, mutations in genes encoding adenylcyclase (*cyaA*) and phosphotransferses (*ptsI*) are known to decrease intracellular levels of cAMP, thereby reducing the expression of *glpT* and *uhpT* and, consequently intracellular fosfomycin levels.[11] Mutations in the gene encoding the drug target MurA, particularly those that confer amino-acid changes in the active site and Cys115 residue have been demonstrated to decrease the susceptibility of the organism by reducing its affinity for fosfomycin.[12-14] However, changes of this nature are uncommon, and have been shown to impair peptidoglycan synthesis and bacterial fitness.[10] Over-expression of *murA* has been found both in mutants selected *in-vitro* and in clinical isolates. It has been suggested this mechanism acts to saturate fosfomycin molecules thereby allowing normal cellular function.[15, 16]

A final and, perhaps emerging mechanism of resistance is the acquisition of enzymes that can inactivate fosfomycin by catalysing the opening of its oxirane ring.[17, 18] Four main fosfomycin resistance enzymes have been described: FosA, a glutathione-S-transferase; FosB a l-cysteine-thiol-transferase; FosC, an ATPase; and FosX, an epoxide-hydrolase. Each catalyses the addition of glutathione, L-cystein, ATP and water respectively to C1 of the oxirane ring of fosfomycin.[19] Genes encoding FosA, B and C are typically found on plasmids when observed in *E. coli*.

Data from multiple studies has shown that exposure to fosfomycin *in-vitro* rapidly selects resistant mutants, at a frequency of 10^−7^-10^−8^.[20, 21] However, mutants selected experimentally are typically physiologically impaired; with decreased growth rates in culture media and urine when compared to wild-type strains.[20] It is also thought that fosfomycin resistant isolates may have a reduced ability to adhere to uroepithelial cells or catheters, and to have a higher sensitivity to polymorphonuclear cells and serum complement killing.[22] Therefore, it has been speculated that despite the rapid development of resistance *in-vitro*, significant biological fitness costs prevent the establishment and propagation of resistant strains *in-vivo*. This may explain the low rates of fosfomycin resistance observed to date in regions where this drug is used.[2, 20]

In Japan, Spain, Germany, Austria, France, Brazil, and South Africa, fosfomycin has been used extensively for >30 years.[23] In these regions a soluble salt form called fosfomycin-tromethamine (typically given as a single 3 g oral dose) is widely used in the treatment of uncomplicated UTIs.[24] Until recently, fosfomycin-trometamol was not distributed or commercially available in the UK; and any products used were imported, typically from Germany and therefore unlicensed. Despite this, the NHS recorded a ten-fold increase in fosfomycin-trometamol prescriptions from 100 to 1000 between 2012 and 2013; [25] [25] and a further increase to 2,400 prescriptions in 2014.[26]

Renewed interest in fosfomycin has been for treatment of MDR organisms causing UTIs where oral therapy choices may be limited. Considering these factors and the possibility of introducing fosfomycin preparations into our formulary, the first aim of this study was to determine the proportion of organisms isolated from UTIs showing resistance to fosfomycin. In doing so the various methods of measuring susceptibility to fosfomycin in the laboratory were compared and their relative merits considered. The second aim was to investigate mechanisms of fosfomycin resistance.

## Materials and Methods

### Bacterial isolates

Between July and August 2014, 2800 urine specimens received as part of standard patient care (over 18 days in total) at Northampton General Hospital, a large 700 bed tertiary hospital in the UK were collected. Subsequent analysis of isolates and susceptibility testing followed the laboratory work-flow and methodologies used for clinical investigation of specimens in this trust. Each was examined for signs of infection using Iris IQSprint microscopy and those meeting conventional clinical criteria were cultured using the MAST Uri®system (n=1000). The susceptibility status of each cultured isolate to fosfomycin was determined using a 96-well ‘breakpoint’ agar plate containing 32 µg/ml fosfomycin supplemented with 25 µg/l of glucose-6-phosphate (G6P) as provided by MAST, and a presumptive species identification was carried out by determining the colour of colonies growing on MAST CUTI chromogenic agar. A total of 62 isolates putatively identified as fosfomycin resistant *E. coli* then had their species confirmed using MALDI-TOF and were retained for further study. *E. coli* J53-2 (NCTC 50167) was used as a fosfomycin susceptible control; *E. coli* NCTC 10418 was used as a quality control for susceptibility testing; and *E. coli* MG1655 (ATCC 700926) was used as a reference strain for genome comparisons.

### Antimicrobial susceptibility testing

The MICs of an extended panel of antimicrobials were determined using the BD Phoenix™ automated microbiology system with AST panel UNMIC-409 as per the manufacturer’s instructions. Fosfomycin MICs were further determined using fosfomycin E-tests^®^ (bioMérieux) and using the agar dilution method adapted from the British Society of Antimicrobial Chemotherapy (BSAC) guidelines.[27]

### Whole genome sequencing (WGS) and post sequencing analysis

Isolates consistently considered resistant by all susceptibility testing methods were genome sequenced by MicrobesNG using an Illumina MiSeq system. Velvet (Version 1.2.10)[28] was used for *de-novo* assembly of the genomes, and Prokka (Version 1.11)[29] used for annotation. Reads were also analysed using the ‘nullarbor’ pipeline (v1.2) using a standard virtual machine on the MRC CLIMB framework. Pan genomes were generated using ‘roary’ (v8.0), SNPs called with ‘snippy’ (v3.0) and antibiotic resistance genes and mutations identified using ‘ARIBA’ (v2.8.1). Trees were visualised with ‘Phandango‘. All packages used default parameters unless stated otherwise. The Centre for Genomic Epidemiology (http://www.genomicepidemiology.org/) provided software for interrogation of genomes for Multi-locus sequence type (MLST), *E. coli* serotype, plasmid replicons and resistance associated genes (ResFinder); the Comprehensive Antibiotic Resistance Database (CARD) was additionally used to seek resistance determinants.[30]

## Results

### Prevalence of fosfomycin resistance in UTI isolates using MAST urisystem

From 1000 UTI culture positive isolates, 20.9% (n=209) were determined to be fosfomycin resistant using ‘breakpoint plates’ on the MAST Uri^®^system, with growth on ≥80% of the culture well indicating an MIC ≥32 μg/ml. Resistance to fosfomycin was observed in a range of bacteria, the largest proportion of which were *E. coli* (n=81). A total of 12% (81/676) of *E. coli* isolates were initially deemed fosfomycin resistant. Of these 81, 62 were confirmed as being *E. coli* by MALDI-TOF analysis. The remaining 19 were found to be either other coliform species or failed to grow on subculture.

### Determination of fosfomycin minimum inhibitory concentrations

#### Fosfomycin MICs using BD Phoenix™

Using an automated micro-broth dilution method (BD Phoenix™) 53/62 *E. coli* isolates (85.5%) were found to have fosfomycin MICs of <16 μg/ml, three isolates had an MIC of 32 μg/ml (4.8%) and six isolates had an MIC of 64 μg/ml (9.7%). Therefore only nine isolates showed concordance with data from the MAST Uri^®^System, and were deemed resistant using BD interpretative software (Epicentre™) with EUCAST breakpoints (≥32 <g/ml). [31]

#### Fosfomycin MICs using E-tests

Due to the discrepancy between micro-broth dilution and breakpoint plate MIC methods, E-tests were used as an alternative method for measuring fosfomycin MICs. Two susceptible control strains, *E. coli* J53-2 and *E. coli* NCTC-10418 grew with definitive zones of inhibition, revealing MICs of 0.25 µg/ml. Similarly, six selected isolates deemed resistant using the MAST Uri^®^system but susceptible using the micro-broth dilution (BD Phoenix™) were found to be sensitive to fosfomycin using E-tests; each growing with a single defined zone of inhibition and MICs ranging from 0.19-0.75 µg/ml (Table 1).

**Table 1:** Fosfomycin minimum inhibitory concentrations and growth characteristics

| Isolate | Fosfomycin MIC (µg/ml) |  |  |
| --- | --- | --- | --- |
|  | MastUri | BD Phoenix | E-test |
| MU721372 | ≥32 | 64 | 512 (24) |
| MU723051 | ≥32 | 64 | 384 |
| MU715908 | ≥32 | 64 | 384 (98) |
| MU720214 | ≥32 | 64 | 384 (48) |
| MU723320 | ≥32 | 64 | 256 |
| MU723292 | ≥32 | 64 | 192 |
| MU720350 | ≥32 | 32 | 256 |
| MU723240 | ≥32 | 32 | 256 (12) |
| MU720142 | ≥32 | 32 | 96 (4) |
| MU723432 | ≥32 | <16 | 0.38 |
| MU724857 | ≥32 | <16 | 0.75 |
| MU719876 | ≥32 | <16 | 0.25 |
| MU724367 | ≥32 | <16 | 0.19 |
| MU725806 | ≥32 | <16 | 0.5 |
| MU725463 | ≥32 | <16 | 0.25 |
| NCTC 10418 | <16 | <16 | 0.25 |
| J53-2 |  |  | 0.25 |

MIC values in brackets represent interpretations of the E-test which include ‘intermediate’ growth within the zone of inhibition

All the isolates deemed resistant by both Mast Uri^®^system and BD Phoenix™ were also categorised as resistant using E-test. Despite agreement of a resistance interpretation between the three methods, there was little concordance between the specific MICs determined by E-tests and the micro-broth dilution method (Table 1). Of note was the difficulty in reading and interpreting E-tests. In each test a small number of single colonies were observed within the clearance zone. As recommended by others who have recorded the same phenomenon,[32] these colonies were excluded from the E-test interpretation. Five isolates had a visible ‘intermediate’ zone of noticeably less dense growth, presenting two possible interpretations. Due to the semi-confluent nature of the growth in these regions they were not included in the zone of inhibition when reading the strips (Table 1).

#### Investigation of fosfomycin MICs using modified agar dilution

To further explore the differing growth phenotypes when using E-tests, a modified agar dilution method was used whereby colonies were streaked on agar containing different concentrations of fosfomycin and their growth observed. For the control organisms and six Phoenix™/E-test determined fosfomycin susceptible organisms, either no growth, or single colony/scanty growth was observed on agar containing a low concentration of fosfomycin (≤ 16 μg/ml). Each of the nine resistant isolates cultured on a low concentration of fosfomycin produced uniform colony morphologies; when grown in the presence of higher concentrations of fosfomycin however each produced a ‘dual colony’ growth phenotype.

### Characterisation of selected E. coli isolates

WGS was used to characterise eight of the consistently fosfomycin resistant isolates and two, randomly selected susceptible isolates. Fosfomycin resistance was present in several different *E. coli* sequence types (6 different STs were seen in the 8 resistant isolates, ST131 was the only ST seen more than once) indicating that resistance was not distributed due to clonal expansion of one strain (Figure 1 and Table 2). The *E. coli* sequence types found in this study include those previously reported as common in UTI isolates in the UK; ST69, 73, 95 and 131.[33]

**Figure 1.**
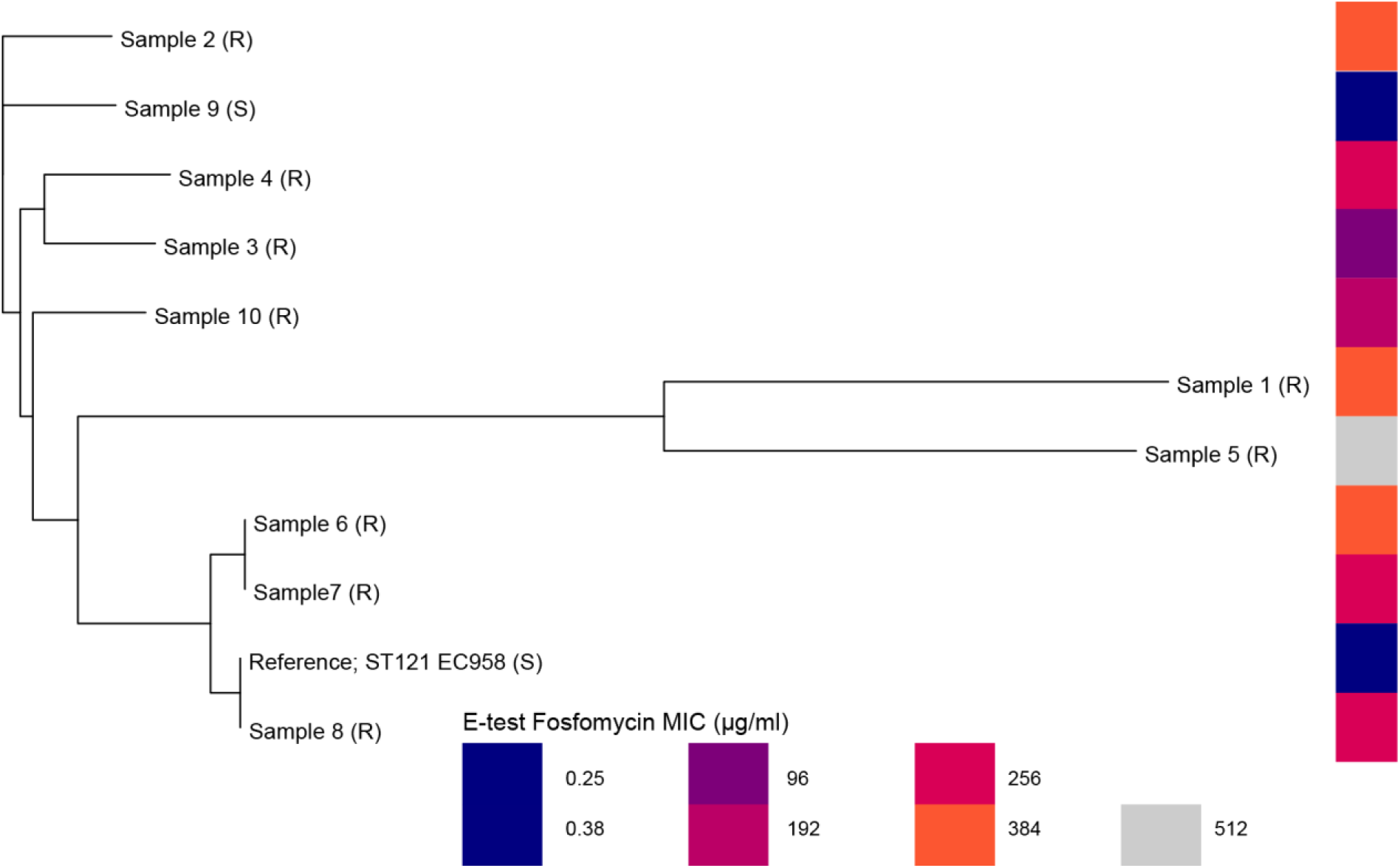
Phylogenetic reconstruction of population structure of the Fosfomycin resistant *E. coli* isolates produced by Roary. (R) and (S) indicate resistant and sensitive isolates respectively. ST 121 strain EC958 was used as a reference.

**Table 2:** Genotypic characterisation of selected *E. coli* isolates

| Isolate | Fosfomycin<br>MIC (Phoenix) | Serotype | ST | Antibiogram<br>(Phoenix) | Resfinder/ CARD:<br>Presence of resistance genes | Plasmid replicons |
| --- | --- | --- | --- | --- | --- | --- |
| MU721372 | 64 µg/ml | O17/O77:<br>H18 | 69 | Fos, Amp, Trim | *bla*<sub>TEM-1B</sub>, *sul2*, *dfrA17*, *aph(6)Ib*,<br>*aph(3')Ib*, | IncFII, IncFIB, Col156, IncQ1 |
| MU723051 | 64 µg/ml | O16:H5 | 131 | Fos, Amp, Cefurox,<br>Gent, Cipro | *bla*<sub>TEM-1B</sub>, *aac(3)-IId*, *gyrA* | IncFII, IncFIB, IncFIA |
| MU715908 | 64 µg/ml | O111:H21 | 40 | Fos | - | - |
| MU720214 | 64 µg/ml | O6:H1 | 73 | Fos, Amp, Trim | *bla*<sub>TEM-1B</sub>, *sul1*, *dfrA5* | IncFIB, Col156 |
| MU723320 | 64 µg/ml | O16:H5 | 131 | Fos, Amp | *bla*<sub>TEM-1B</sub> | IncFII, IncFIB, Col156 |
| MU720350 | 32 µg/ml | O75:H5 | 550 | Fos | - | IncFII, IncFIB, IncX1, Col156,<br>Col |
| MU723240 | 32 µg/ml | -:H4 | 131 | Fos, Amp, Trim | *bla*<sub>TEM-1B</sub>, *sul1*, *dfrA17*, *aadA5*, | IncFII, IncFIA |
| MU720142 | 32 µg/ml | O6:H31 | 127 | Fos, Amp, Trim | *bla*<sub>TEM-1B</sub>, *sul2*, *dfrA14*, *aph(3')Ib*,<br>*aph(6)Ib* | IncFII, IncFIB, IncB/O/Z/K,<br>Col156 |
| MU723432 | <16 µg/ml | O83:H33 | 567 | Amp, Trim | *bla*<sub>TEM-1B</sub>, *sul2*, *dfrA8*, *dfrA14*,<br>*strB*, *aph(3')Ib*, *aph(6)Ib*, | IncFII, IncFIB, IncFII(pCoo) |
| MU724857 | <16 µg/ml | O25:H4 | 95 | Amp, Coamox,<br>PipTaz | *bla*<sub>TEM-1B</sub> | IncFII, ColpVC, IncFIB,<br>IncB/O/Z/K |

Fos, fosfomycin; Amp, ampicillin; Trim, trimethoprim; Coamox, coamoxiclav; Cefurox, cefuroxime; Cipro, ciprofloxacin; Gent, gentamicin; PipTaz, Tazocin

Each of the ten isolates were further characterised by investigating their antibiogram, determined from their susceptibility profile to antimicrobials used in the treatment of UTIs; and by interrogating WGS for genes and mutations known to confer antimicrobial resistance (Table 2). Ampicillin resistance was detected in 8/10 isolates, accompanied with the *in-silico* detection of *bla*_TEM-1B_. Trimethoprim resistance in 5/10 isolates corresponded with the detection of a *dfrA* gene and with either *sul1* or *sul2*. Aminoglycoside resistance genes were identified in five of the isolates where in one ST131 isolate, the presence of *aac(3)-IId* and a *gyrA* mutation corresponded to gentamicin and ciprofloxacin resistance respectively.

The *in-silico* analysis also showed the presence of many common *Enterobacteriaceae* plasmid replicons including those of incompatibility group, IncF, IncQ, IncX1, IncB/O/K/Z and plasmids from the group Col and Col156. Using CARD, Resfinder and manual searches, no fos-like genes were detected in any of the strains, suggesting an absence of known plasmid based transferrable fosfomycin resistance genes in the resistant isolates.

### Amino-acid variation in proteins associated with fosfomycin resistance

For each of the ten sequenced isolates, amino-acid changes or mutations in known fosfomycin resistance genes *murA*, *glpT*, *uhpT*, *uhpA*, *ptsI* and *cyaA* were identified from the WGS using *E. coli* MG1655 as a reference (Table 3). No *murA* changes were identified in any of the fosfomycin resistant isolates, a single substitution of Val389Ile was found in susceptible isolate, MU723432.

**Table 3:**
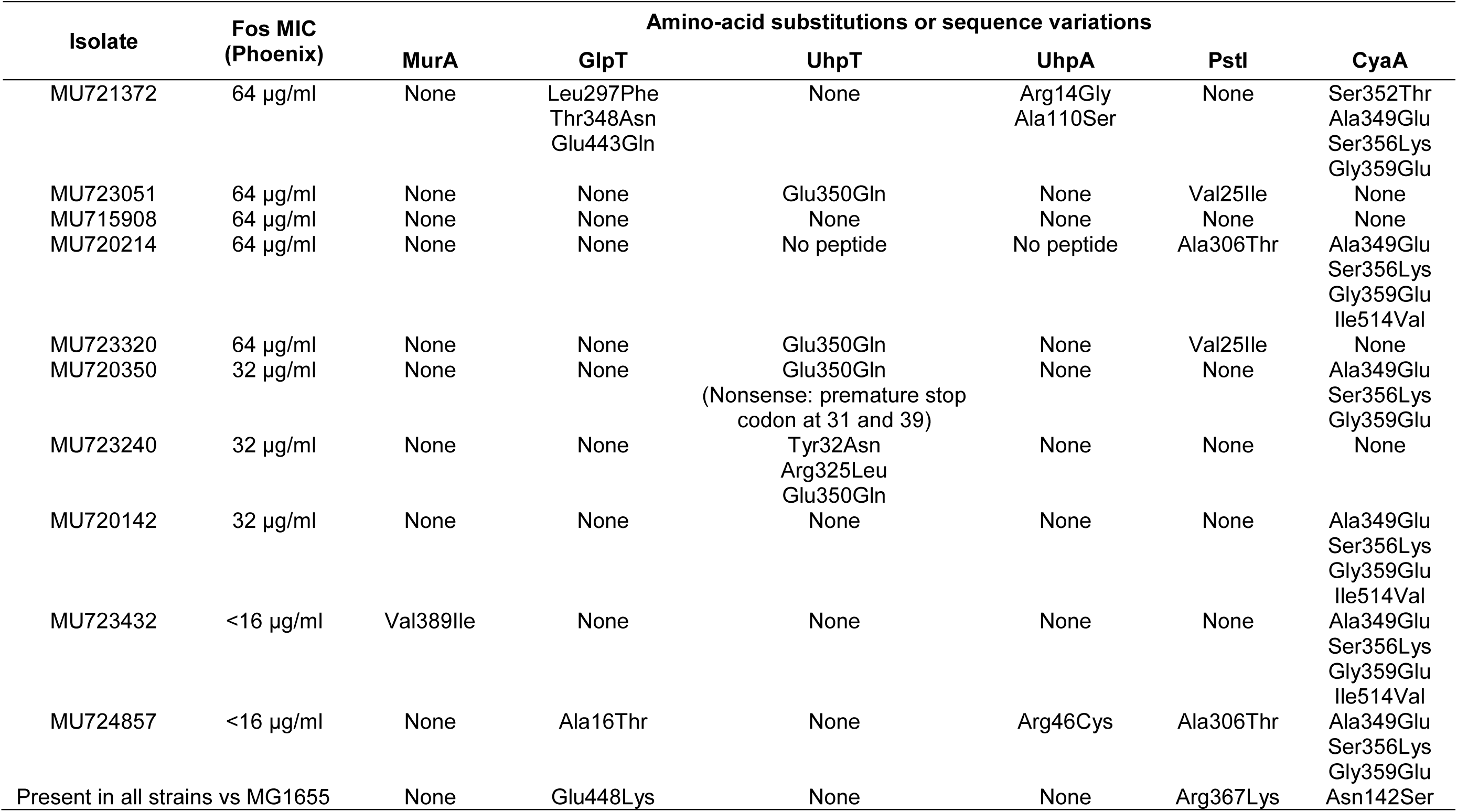
Fosfomycin-associated mutations found in resistant *E. coli* isolates

All sequenced isolates were found to have a Glu448Lys change in GlpT when compared to MG1566. Fosfomycin resistant isolate MU721372 had an additional three substitutions of Leu297Phe, Thr348Asn, Glu443Gln, however susceptible isolate MU724857 also had a second GlpT change of Ala16Thr.

No amino-acid changes in the sequence of UhpT were identified in fosfomycin susceptible isolates; however, 5/8 resistant isolates had changes in this protein. In MU720214, both *uhpT* and *uhpA* were completely absent. Comparative analysis against other *E. coli* genomes showed the presence of a phage integrase gene adjacent to the *uhpT-uhpA* region within the assembled contig, suggestive of a deletion event. Isolate MU720350 had two amino-acid changes at positions 31 and 39 predicted to confer premature stop codons leading to a truncated protein; four strains had a Glu350Gln amino-acid substitution; and MU723240 had additional substitutions of Tyr32Asn and Arg325Leu.

Only three isolates had changes in the *uhpA* gene, a deletion in MU720214, an Arg46Cys substitution in susceptible isolate MU724857, and substitutions Arg14Gly and Ala110Ser in fosfomycin resistant isolate MU721372 (Table 3).

When examining genes that affect levels of intracellular cAMP, all the isolates had the substitution of Arg367Lys in PtsL and Asn142Ser in CyaA when compared to MG1655; both changes are well represented in many *E. coli*. Two further substitutions were identified in PtsL, Val25Ile in two of the resistant isolates (MU723051 and MU723320) and Ala306Thr in one resistant (MU720214) and one susceptible *E. coli* (MU724857). The amino-acid sequences of CyaA in each isolate fell broadly into two groups, those with a single Asn142Ser change when compared to MG1655 (n=4), and those with ≥3 additional amino-acid substitutions (Ser352Thr, Ala349Glu, Ser356Lys, Gly359Glu and Ile514Val) (n=5 Table3) both containing susceptible and resistant isolates. These amino-acid substitutions appeared to correlate more closely with sequence type than with fosfomycin susceptibility status and were found commonly in other *E. coli* strains.

## Discussion

To determine the prevalence of fosfomycin resistance from a panel of UTI isolates, different methods to distinguish susceptible and non-susceptible isolates in a clinical laboratory were explored. These followed the order which would be used clinically in this setting. Use of ‘breakpoint’ plates on the MAST Uri^®^system for high throughput screening determined the prevalence of resistance (MIC ≥32 μg/ml) in *E. coli* isolates as 12%; a rate significantly higher than previously documented.[34-36] However, on further examination using automated micro-broth dilution, only nine of these isolates were resistant (MIC ≥32 μg/ml). Furthermore, if CSLI guidelines had been applied none of the isolates would be deemed resistant, as each had an MIC below the breakpoint according to this scheme (S≤64, I=128 and R ≥256 μg/ml). [37][37] Susceptibility interpretations from the E-test method corroborated the findings from micro-broth dilution, concordantly differentiating isolates deemed fosfomycin susceptible and resistance. Therefore, both these methods agree that only 1.3% of *E. coli* within the study should be regarded as fosfomycin resistant using current definitions; a prevalence more in line with findings of previous studies both globally and within the UK.[38, 39] The high prevalence of resistance recorded by the MAST Uri^®^system reflects a large number of false positive results (53/62) given the interpretive criteria followed. Whilst changes to fosfomycin susceptibility can occur relatively rapidly *in-vitro* it is infeasible that a significant number of isolates initially identified as resistant would have reverted to susceptibility in the time window of the laboratory investigations. In the collection period, fosfomycin was not used in the trust or by community pharmacists in this area, thereforepatient exposure to the drug is likely to have been low, and a 1.3% rate of resistance is likely to reflect spontaneous mutants which are in the wider population of *E. coli.* Given the reports of fosfomycin resistance incurring a significant fitness cost this level may be higher than expected given the probable lack of direct selection in this population.

In this study, micro-broth dilution was the most useful method for differentiating resistant and susceptible isolates, others have found disc-diffusion assays such as those described by Lu *et al.* [40] to be the most reliable; despite reports of single colony generation within the zone of inhibition.[32] A beneficial next step might be to directly compare micro-broth dilution and disc-diffusion assays to determine the most robust and practical method for determining fosfomycin susceptibilities within a clinical laboratory setting. Interpretation of E-tests was obfuscated by an intermediate zone of growth, resembling in appearance a ‘small’ colony phenotype observed at higher concentrations of fosfomycin when isolates were streaked onto plates. A similar ‘dual colony’ phenomenon in the presence of fosfomycin has been described previously by Tsuruoka *et al.* who reported differences in growth and carbohydrate uptake between colony types.[21] In the present study, these distinct phenotypes were found to be transient and inconsistent, large and small colonies going on after passage to produce daughter colonies of both phenotypes in the presence of higher concentrations of fosfomycin (data not shown) further hindering interpretation of susceptibility testing.

*In-silico* MLST and whole genome comparison of the fosfomycin resistant *E. coli* showed that the isolates were of diverse sequence-types, and that resistance and plasmid profiles differed in each isolate. Therefore, resistance had not disseminated in this population due to expansion of one clone. Examination of the mechanisms of resistance found no evidence for mobile elements being involved in fosfomycin resistance, the absence of any plasmid located *fos* genes suggests that resistance in these *E. coli* was due to chromosomal mutations. When examining sequences of genes known to contribute to fosfomycin resistance, no two isolates had the same set of substitutions or mutations. As in other studies, changes in GlpT and UhpT/UhpA transport systems responsible for uptake of fosfomycin were the most commonly identified; with 6/8 resistant organisms possessing amino-acid changes or deletions within these systems that were absent in the susceptible strains. This included the complete deletion of the *uhpT*/*uhpA* region; location of a premature stop codon predicted to lead to a truncated UhpT protein; and the commonly reported UhpT substitution Glu350Gln;[14, 41] all speculated to result in reduced uptake of fosfomycin. Substitutions in GlpT were less common in this study than other recent reports, only a single isolate (MU721372) accumulating many changes in this region. Of note is the Glu448Lys substitution, identified previously in other fosfomycin resistant isolates.[14] This change was identified in all the sequenced isolates when compared to MG1655, including those deemed susceptible, but was not found during a search of an extended panel of sequenced *E. coli* submitted to Genbank. This suggests that either this substitution does not confer resistance to fosfomycin, contradicting speculation by others;[14] or that it acts to reduce susceptibility, perhaps below our defined breakpoints in the absence of other changes within the protein. It may be that low-level changes to susceptibility account for why some isolates were deemed resistant using screening with the MAST Uri^®^system, whilst remaining sensitive using other testing methods.

Only a single substitution (Val389Ile) was identified in MurA within the sequence of one of the susceptible isolates. Although the modification has been reported by others in fosfomycin resistant isolates,[41] its location outside the active site of this enzyme means its role in resistance is ambiguous. The role of changes in CyaA and PtsI proteins in this study is less clear. The amino-acid sequence of CyaA appeared to divide into two groups both with substitutions which can be found in other fosfomycin susceptible *E. coli*. This suggests that these changes may be unrelated to fosfomycin susceptibility, but may correspond to the *E. coli* phylogeny. While many of the substitutions identified in this study have previously been linked to fosfomycin resistance by others, our detection of amino acid changes in both susceptible and non-susceptible strains raises doubts regarding their contribution to fosfomycin resistance.

The use of fosfomycin for treatment of UTIs and other infections is likely to increase. In this study, the prevalence of fosfomycin resistance in *E. coli* isolated from UTIs was found to be relatively low and resistant isolates were divergent. The identification of chromosomal based changes in genes associated with fosfomycin susceptibility, and the absence of *fos* genes on conjugative plasmids indicates that resistance in these isolates was not transferrable, and that co-location with other resistance genes did not appear to lead to co-selection. Therefore, in this setting fosfomycin remains a useful agent in the treatment of UTIs, equipping us with an extra option for hard to treat UTIs and providing an alternative to drugs such as carbapenems which may drive selection of resistant organisms further. Current methods to identify fosfomycin resistant *E. coli* isolates in urine can give very different results, there is a need for more consistency to accurately define real rates of resistance which is important in monitoring any evolution of resistance as fosfomycin use is likely to increase.

## Acknowledgements

This work was carried out with the support of the staff in the microbiology department at Northampton General Hospital

## Transparency declarations

None to declare.

